# A Novel No-Wash (Homogenous) High-Throughput Toxicity Assay on Primary Cells

**DOI:** 10.1101/2025.04.21.649658

**Authors:** Yan Cheng, Victoria Yang, Zixuan Zhu, Grace Yang, Jianming Lu

## Abstract

Mitochondrial membrane potential (MMP) is a critical marker of mitochondrial and, therefore, cellular function. However, current dyes used for MMP measurement possess significant limitations: nonspecific targeting, long incubation times, potential mitochondrial toxicity, poor water solubility, and numerous wash steps. Using a novel fluorescent dye—mitochondrial membrane potential indicator (m-MPI)—in combination with a masking dye, we overcame said limitations and successfully measured MMP changes on different cell lines, as well as four different human primary cells. This no-wash assay has been miniaturized into a 384-well format. It is robust with Z’ value > 0.5 on both HepG2 and HeLa cells and can be used for high throughput screening. The reagents in the m-MPI homogeneous assay kit are not toxic to cells, as was demonstrated on HepG2 and HeLa cell lines using Dexorgen’s EnerCount ATP assay. Using this assay, we conducted a mini screening on human hepatocytes against the Library of Pharmacologically Active Compounds (LOPAC) (Sigma). The resulting hits were then confirmed with follow-up studies.

## INTRODUCTION

Mitochondria are double membrane-bound organelles found in most eukaryotic cells and primarily function to produce ATP via respiration. Mitochondria play an important role in maintaining cellular homeostasis and are involved in a variety of critical cellular processes, including apoptosis (1).

Chemically induced mitochondrial toxicity is a critical evaluator of chemical cytotoxicity in eukaryotic cells because of the central role mitochondria plays in energy production, apoptosis, and necrosis regulation (2,3). Such toxic effects on mitochondria are indicated by changes in mitochondrial membrane potential (MMP). Thus, current toxicity assays have been developed using membrane-permeable, fluorescent lipophilic cations to determine the integrity of MMP. Although these assays are important components of a toxicological screen, the dyes used have their limitations. They can accumulate in membrane components other than mitochondria (4) or require an extended period to reach mitochondrial membrane equilibrium, especially when the cellular accumulation of the dye is reduced by the multi-drug resistance pump (5). Other assays may also exert toxic effects on mitochondria (6,7).

Examples of fluorescent dyes used to measure MMP include nonylacridine orange (NAO), safranine O, rhodamine 123, chloromethyl-tetramethyl-rosamine, and tetramethylrhodamine methyl and ethyl esters (TMRM and TMRE). Two popular dyes are 3,3’-Dihexiloxadicarbocyanine iodide [DiOC6(3)] and the cyanine dye 5,5’,6,6’-tetrachloro-1,1’,3,3’tetraethyl benzimidazolyl carbocyanine iodide (JC-1) (8). DiOC6(3) works effectively in isolated mitochondria, but not in whole cells, because it detects both plasma and MMP changes (8). JC-1 (9) has an advantage over rhodamines and other carbocyanines because it is more selective to mitochondria. The time required for JC-1 to reach equilibrium is relatively short, and it has minimal toxic effects on the mitochondrial electron transport chain (10). JC-1 aggregates and accumulates in the mitochondria of healthy cells but remains in the cytoplasm as a monomer in less healthy cells with lower MMP. It is relatively specific and sensitive with a low background (11). However, the most significant limitation of JC-1 is its poor water solubility: JC-1 begins to precipitate in aqueous buffers at concentrations as low as 1 μM due to its hydrophobic nature. This limitation makes it very difficult to load consistent amounts of JC-1 into cells, resulting in large experimental variation. JC-1 is also not suitable for all cell types. For instance, it can be used to measure MMP changes in HeLa cells but not in HepG2, CHO, or primary rat hepatocytes due to its low signal-to-background window (7). Additionally, although JC-1 is a very specific dye, it is not particularly sensitive in terms of changes in MMP (12).

To overcome the limitations of the MMP assays described above, we have developed a no-wash, high-throughput, cell-based assay using the novel dye m-MPI that has been tested on multiple different cell lines and primary cells (7,13).

m-MPI is a modified MMP sensor that is based on JC-1 but is more water-soluble. In healthy cells, mitochondria are polarized and m-MPI accumulates in the mitochondria as aggregates with red fluorescence (emission at 590 nm). In cells with lower MMP, m-MPI remains in the cytoplasm as monomers that emit green fluorescence (emission at 535 nm). The dye undergoes a change in fluorescence emission from green to red when the MMP increases, or vice versa, and is a reversible process (13). A masking dye is also used to cover the fluorescence from m-MPI located outside of the cells (Fig. 1). The green-to-red fluorescence ratio relies only on the membrane potential, not the mitochondrial size, shape, or density, and can be used to determine the mitochondrial function of the cells (10) (Fig. 1). *Sakamuru*, et al (13) compared m-MPI to JC-1, rhodamine123, and TMRE with three known electron transport chain inhibitors within the mitochondria—FCCP, rotenone, and antimycin A—in HepG2 cells and found that the m-MPI dye is superior to all three other dyes for the MMP assay.

**Fig. 1.**
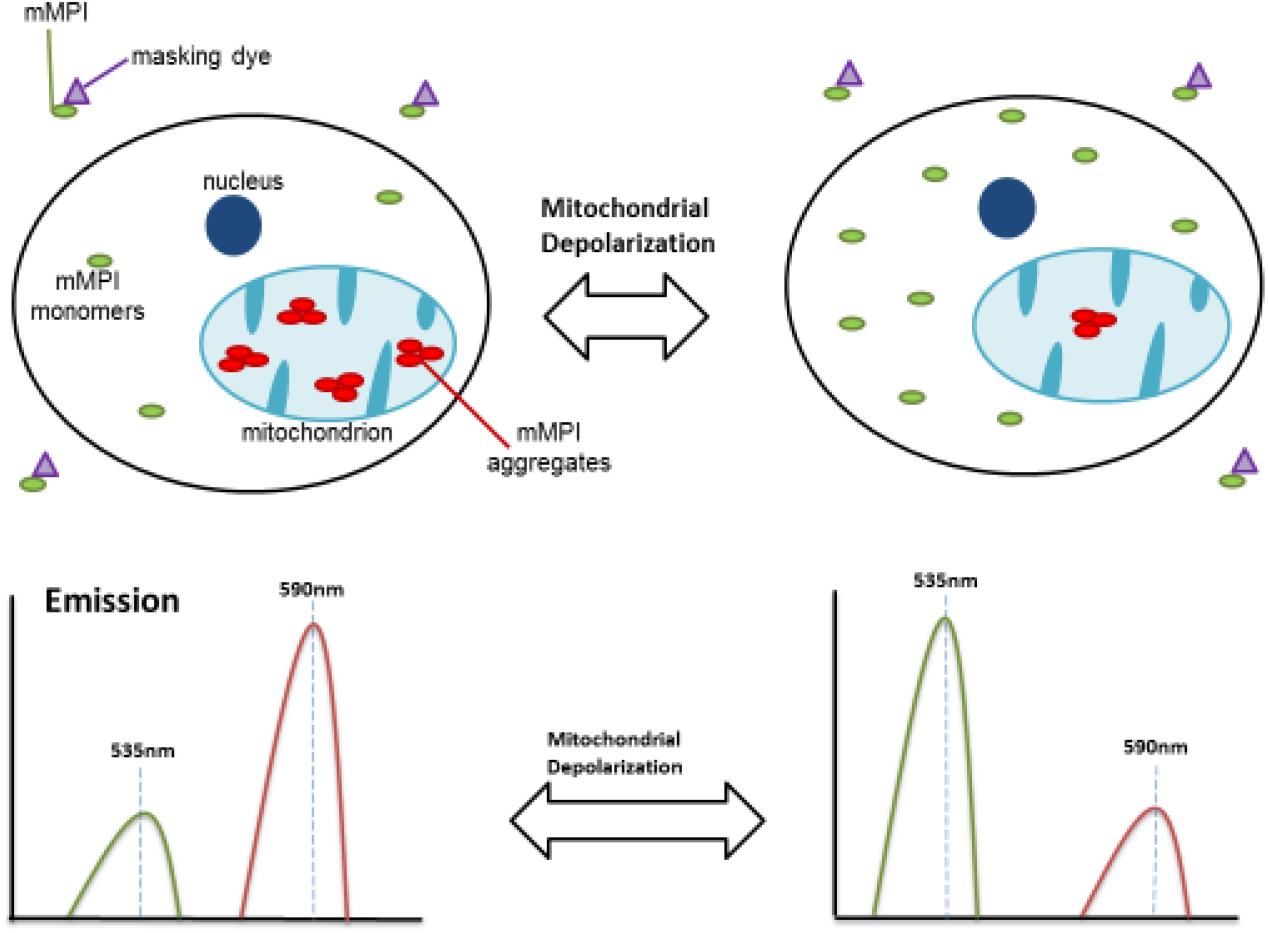
Schematic illustration of m-MPI dye and masking dye in mitochondria to measure mitochondrial membrane polarization.

To reiterate, this report details the development of a no-wash (homogenous) MMP assay that can be applied to many different cell lines and primary cells.

## MATERIALS AND METHODS

### 1. Cell Culture

HepG2 (Hepatocellular carcinoma), CHO-K1, HEK293, and HeLa cells were purchased from the American Type Culture Collection (ATCC, Manassas, VA, USA). HepG2, HEK293, and HeLa cells were cultured in Dulbecco’s Modified Eagle’s Medium (DMEM) (Invitrogen, CA, USA) supplemented with 10% FBS (Sigma-Aldrich, St. Louis, MO, USA), 50 U/ml penicillin, and 50 μg/ml streptomycin (Invitrogen, CA, USA). The cells were maintained at 37°C under a humidified atmosphere and 5% CO_2_. CHO-K1 cells were cultured in Dulbecco’s Modified Eagle Medium: Nutrient Mixture F-12 (DMEM/F12) (Invitrogen, CA, USA) supplemented with 10% FBS (Sigma-Aldrich, St. Louis, MO, USA), 50 U/ml penicillin, and 50 μg/ml streptomycin (Invitrogen, CA, USA).

Human hepatocytes were purchased from ZenBio as frozen vials. The cells were thawed in a 37 ° C water bath and re-suspended in the Plating Medium. Human hepatocytes were plated on 384-well collagen-coated plates at the density of 9K/well (in 20 μl Plating Medium) and incubated in a 37 C incubator.

Human adult stem cells were purchased from Zen-Bio, Inc. (Research Triangle Park, NC). The cells were maintained and proliferated in Preadipocyte Medium (PM-1). 2.3K human adult stem cells in 25 μl of PM-1 were plated into each well of a 384-well plate and grown in a 37°C CO_2_ incubator for 24 hrs. The PM-1 was then removed and replaced with 25 μl of Adipocyte Difference Medium (DM-2). The cells were incubated in a CO_2_ incubator for seven days. 25 μl of DM-2 was removed and replaced with 25 μl of Adipocyte Medium (AM-1), after which the human adult stem cells were incubated for another seven days and differentiated into adipocytes.

Human epidermal melanocytes, neonatal, lightly pigmented donor (HEMn-LP) were purchased from Life Technologies (Grand Island, NY). The cells were maintained and proliferated according to the manufacturer’s protocol.

Human skeletal myoblasts (HSkM-L) were purchased from Life Technologies (Grand Island, NY). The cells were thawed from a cryo-vial and plated on a 384-well plate in 50 μl of Differentiation Medium with a density of 12K cells/well. The cells were incubated in a CO_2_ incubator for 48 hours to enable rapid differentiation.

### 2. Reagents

A library of pharmacologically active compounds (LOPAC), containing 1280 compounds with known pharmacological actives, was purchased from Sigma (St. Louis, MO). All the chemicals for confirmation assay except tyrphostin 47 were purchased from Sigma (St. Louis, MO). Tyrphostin 47 was purchased from Cayman Chemical (Ann Arbor, MI, USA). All other chemicals were purchased from Sigma unless the chemical providers are otherwise specified.

### 3. Cell-based m-MPI assay on cell lines

12K of HEK-293, 6K of HepG2, 8K of HeLa, or 10K of CHO-k1 cells in 20 μl culture medium were plated into each well of a 384-well black-clear plate. Different concentrations of the testing compounds (2X final concentrations) were prepared in 1X HBSS. An equal volume of the solutions was added into each well and incubated with the cells for 30 min. 3X homogeneous m-MPI solution was prepared according to manufacturer protocol. Equal volumes of the above solutions (3X) were then applied to the cells treated with the compounds. The plate was incubated at 37°C for another 30 min. The signal was recorded via a SpectraMax Gemini EM plate reader (Molecular Devices, Sunnyvale, CA).

### 4. Cell-based m-MPI assay on primary cells

Human hepatocytes from ZenBio (same batch as in the previous report) were placed on 384-well collagen-coated plates at a density of 9K/well (in 25 μl Plating Medium) and incubated at 37°C. On the second day, the plates were removed from the CO_2_ incubator. The medium was removed. 10 μl of fresh Plating Medium was added into each well. Different concentrations of the compounds (2X) were prepared in 1X HBSS. 10 μl of the compound solution was added into each well of the plates. The plates were incubated at 37 °C for the time indicated. Afterward, 3X m-MPI dye solution was prepared in Dexorgen’s proprietary enhancer solution. 10 μl of the 3X dye solution was added into each well and incubated with the cells at 37° C for another 30 min.

The 384-well plate containing differentiated adipocytes was taken out from the incubator and the medium was removed. 20 μl of AM-1 was added into each well of the plate. Different concentrations of the compounds (2X) were prepared in 1X HBSS. 20 μl of the compound solution was added and incubated with the cells for 60 min at 37° C. 20 μl of the 3X m-MPI dye solution was added into each well and incubated with the cells at 37 °C for another 30 min. Human epidermal melanocytes, neonatal, lightly pigmented donor (HEMn-LP) were plated into a 384-well plate at a density of 20K cells/well and incubated in a CO2 incubator for 2 days. On the third day, the plate was then taken out from the incubator and the medium was removed and replaced with 20μl of fresh culture medium. Different concentrations of the compounds (2X) were prepared in 1X HBSS. 20 μl of the compound solution was added and incubated with the cells for 60 min at 37 C. 20 μl of the 3X m-MPI dye solution was added into each well and incubated with the cells at 37° C for another 30 min.

Human skeletal myoblasts (HSkM-L) were thawed from a cryo-vial and plated on a 384-well plate in 50 μl of Differentiation Medium with a density of 12K cells/well. The cells were incubated in a CO_2_ incubator for 48 hours to enable rapid differentiation. Afterward, the plate was taken out from the incubator and the medium was removed. 20 μl of fresh Differentiation Medium was added into each well of the plate. Different concentrations of the compounds (2X) were prepared in 1X HBSS. 20 μl of the compound solution was added and incubated with the cells for 60 min at 37 C. 20 μl of the 3X m-MPI dye solution was added into each well and incubated with the cells at 37 °C for another 30 min. Fluorescence intensities (485 nm excitation, and 535 and 590 nm emissions) were measured using a SpectraMax Gemini EM plate reader (Molecular Devices, Sunnyvale, CA).

### 5. ATP assay on HepG2 and Hela cells

24K of HepG2 and 32K of Hela cells were plated on 96-well plates in 80 ul of culture medium. On the second day, an equal volume of 2X m-MPI homogenous dye solution was added into each well and incubated for 30 min, 1 hour, 2 hours, and 4 hours. The ATP level was then determined with Dexorgen’s EnerCount ATP assay kit (Dexorgen, Inc. Rockville, MD).

### 6. Z’ Value calculation

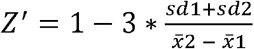

Where:

X2: Average fluorescence intensity from group 2 (cells treated with the compound);

X1: Average fluorescence intensity from group 1 (untreated/control cells); Sd2: standard deviation from group 2 (treated cells);

Sd1: standard deviation from group 1 (control cells)

## RESULTS

### 1. Development of the homogenous m-MPI essay on the 384-well plate

12K of HEK-293 cells were plated into each well of a 384-well plate in 20 μl culture medium. On the second day, 20 μl of different concentrations of FCCP prepared in 1X HBSS (2X of the final concentrations) was added into each well and incubated with the cells for 30 min. At the same time, 3X m-MPI dye or 3X m-MPI in combination with masking dye was prepared in Dexorgen’s proprietary enhancer solution. 20 μl of above solutions (3X) was then applied to the cells treated with FCCP. The plate was incubated at 37°C for another 30 min. The cells treated with 3X m-MPI dye alone (no masking dye) were washed once with Dexorgen’s proprietary enhancer solution (wash protocol). The cells treated with 3X m-MPI dye in combination with masking dye were kept as was (homogenous assay). The plate was placed on a SpectraMax Gemini EM and the fluorescent intensities of both the green (Excitation: 485 nm; Emission: 530 nm; Cutoff: 515 nm) and the red (Excitation: 485 nm; Emission: 590 nm; Cutoff: 570 nm) channels were recorded using the endpoint mode. The ratio of green-to-red signal was plotted (Fig. 2A).

**Fig. 2.**
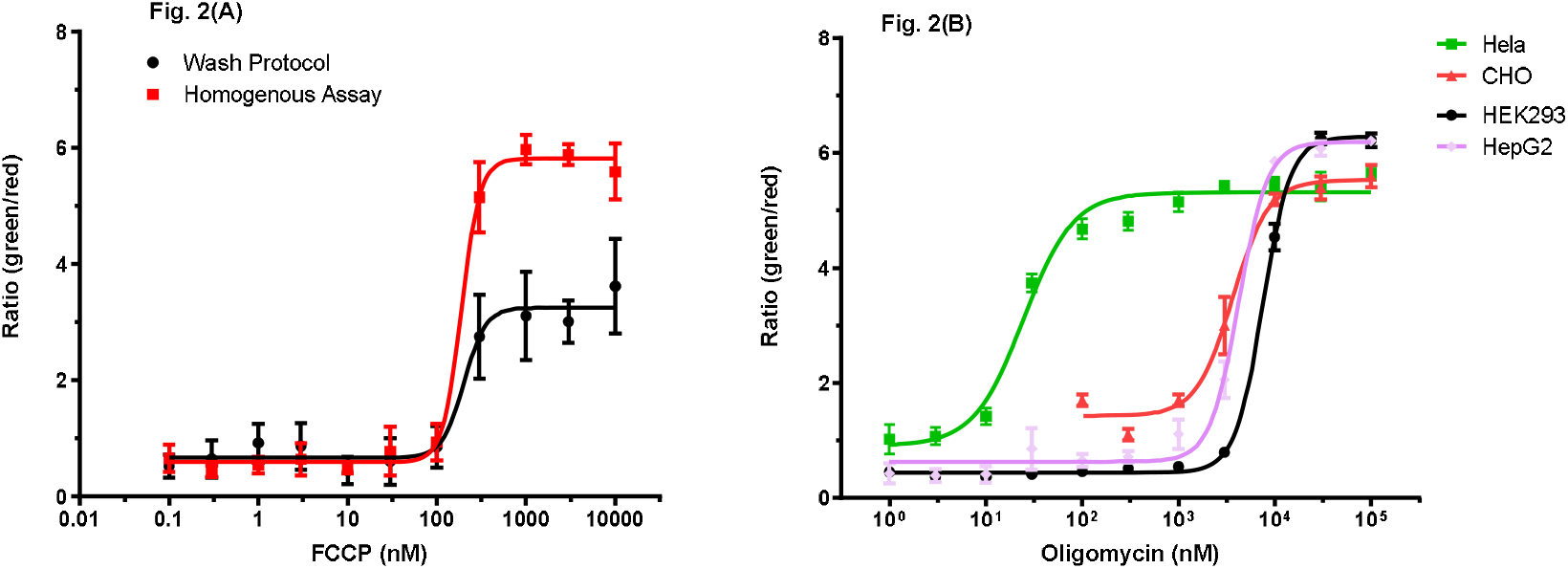
Homogeneous m-MPI assay on 384-well plate using different cell lines. (A) Development of the no-wash assay using HEK-293 cells; (B) Results of homogenous assay using different cell lines

Afterwards, four different cell lines were tested. 6K of HepG2 cells, 10K of CHO cells, 12K of HEK293 cells, and 8K of Hela cells were plated on 384-well plates (black-clear). On the second day, the cell plates were taken out from the CO_2_ incubator. Different concentrations of oligomycin (2X) were prepared in 1X HBSS. 20 μl of the compound solutions was added into each well and incubated with the cells for 30 min. 3X dye solution containing 3X m-MPI plus masking dye was prepared in Dexorgen’s proprietary enhancer solution. 20 μl of the 3X dye solution was added into each well and incubated with the cells for another 30 min. The plate was placed on a SpectraMax Gemini EM and the fluorescent intensities of both the green (Excitation: 485 nm; Emission: 530 nm; Cutoff: 515 nm) and the red (Excitation: 485 nm; Emission: 590 nm; Cutoff: 570 nm) channels were recorded using the endpoint mode. The ratio of green-to-red signal was plotted (Fig. 2B).

### 2. m-MPI assay on Hela cells with different cytotoxic compounds

Twelve mitochondrial toxic compounds were tested on HeLa cells using the m-MPI homogenous assay. The compounds were diluted in 1X HBBS and the assay was performed as described above. The results indicate that the assay is sensitive to those different cytotoxic compounds (Fig. 3).

**Fig. 3.**
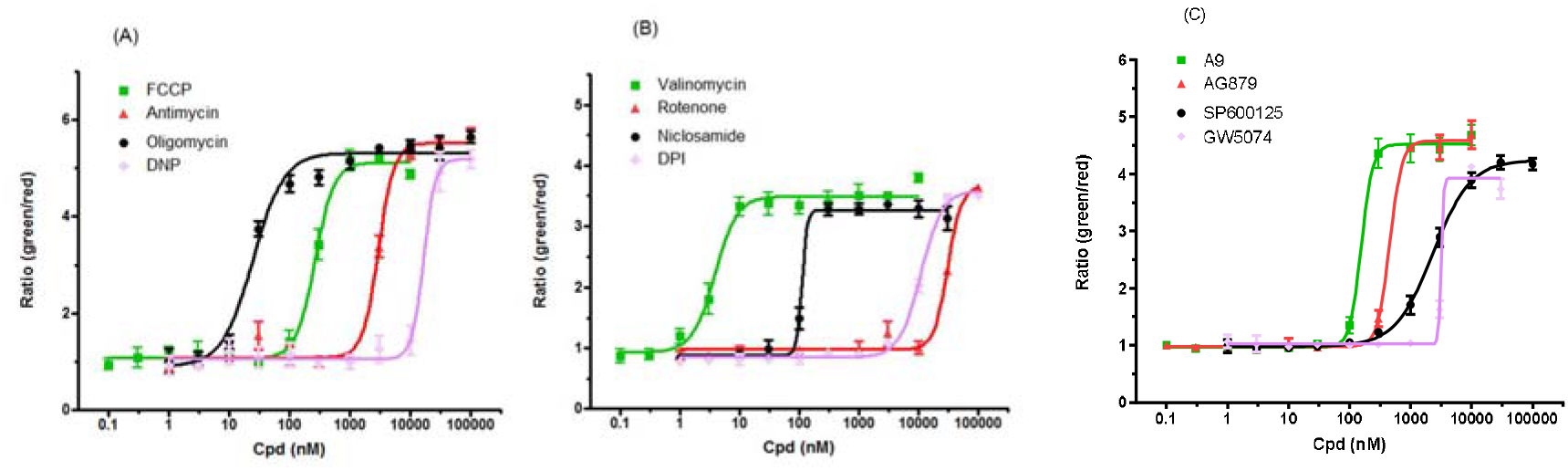
Homogeneous assay with different compounds on HeLa cells. (A): assay using FCCP, antimycin, oligomycin, and DNP; (B): valinomycin, rotenone, niclosamine, and DPI; (C): A9, AG879, SP600125, and GW5074

The 12 compounds were also tested on HEK293, CHO-K1, and HEPG2 cells. The homogenous assay also worked on these different cell lines (data not shown).

### 3. Assay Quality of m-MPI Dye for HTS

6K of HepG2 cells were plated on 384-well plates (black-clear) in 20 μl culture medium. On the second day, the cell plates were taken out from the CO_2_ incubator. 20 μM FCCP was prepared in 1X HBSS. 20 μl of the FCCP solution was added into each well of the right half of the plate and 20 μl of the 1X HBSS was added into each well of the left half of the plate. The cell plate was incubated in a 37°C incubator for 30 min. 3X dye solution containing 3X m-MPI plus masking dye was prepared in Dexorgen’s proprietary enhancer solution. 20 μl of the 3X dye solution was added into each well and incubated with the cells for another 30 min. The plate was placed on a SpectraMax Gemini EM and the fluorescent intensities of both the green (Excitation: 485 nm; Emission: 530 nm; Cutoff: 515 nm) and the red (Excitation: 485 nm; Emission: 590 nm; Cutoff: 570 nm) channels were recorded using the endpoint mode. The ratio of green-to-red signal was calculated. The Z’ value was determined as described in. It is 0.819 on HepG2 cells.

**Table. 1a.** The ratio of Green/Red signal of each well (HepG2 cells) treated with 1XHBSS or 10 μM FCCP. Z’=0.819.

| 1X HBSS | | | | | | | | | | | | 10 μM FCCP | | | | | | | | | | | |
| --- | --- | --- | --- | --- | --- | --- | --- | --- | --- | --- | --- | --- | --- | --- | --- | --- | --- | --- | --- | --- | --- | --- | --- |
| 0.483 | 0.62 | 0.579 | 0.737 | 0.504 | 0.682 | 0.552 | 0.769 | 0.503 | 0.743 | 0.577 | 0.67 | 4.729 | 4.647 | 4.615 | 4.848 | 4.732 | 4.412 | 4.899 | 4.674 | 5.016 | 4.775 | 4.611 | 4.716 |
| 0.608 | 0.629 | 0.643 | 0.641 | 0.635 | 0.584 | 0.634 | 0.615 | 0.647 | 0.592 | 0.631 | 0.599 | 4.683 | 4.809 | 4.714 | 4.264 | 4.954 | 4.598 | 4.8 | 4.521 | 4.656 | 4.878 | 4.536 | 4.713 |
| 0.536 | 0.658 | 0.682 | 0.663 | 0.521 | 0.687 | 0.646 | 0.681 | 0.49 | 0.614 | 0.59 | 0.628 | 4.765 | 4.729 | 4.792 | 4.647 | 4.451 | 4.808 | 4.676 | 4.701 | 4.189 | 4.553 | 4.278 | 4.623 |
| 0.74 | 0.919 | 0.594 | 0.717 | 0.617 | 0.714 | 0.61 | 0.664 | 0.638 | 0.681 | 0.565 | 0.641 | 4.517 | 4.635 | 4.625 | 4.666 | 4.877 | 4.938 | 4.736 | 4.722 | 4.678 | 4.432 | 4.364 | 4.367 |
| 0.544 | 0.774 | 0.618 | 0.695 | 0.518 | 0.614 | 0.585 | 0.718 | 0.682 | 0.804 | 0.6 | 0.699 | 4.912 | 4.882 | 4.856 | 4.934 | 5.01 | 4.904 | 4.612 | 4.616 | 4.671 | 4.526 | 4.494 | 4.432 |
| 0.59 | 0.734 | 0.602 | 0.602 | 0.575 | 0.585 | 0.55 | 0.565 | 0.597 | 0.582 | 0.615 | 0.593 | 4.809 | 4.646 | 4.867 | 4.773 | 4.45 | 4.935 | 4.824 | 4.689 | 4.678 | 4.579 | 4.71 | 4.687 |
| 0.527 | 0.642 | 0.598 | 0.6 | 0.529 | 0.624 | 0.596 | 0.602 | 0.546 | 0.598 | 0.741 | 0.553 | 4.751 | 4.503 | 4.633 | 4.71 | 4.956 | 4.695 | 4.69 | 4.675 | 4.563 | 4.729 | 4.527 | 4.396 |
| 0.548 | 0.67 | 0.631 | 0.654 | 0.582 | 0.629 | 0.665 | 0.881 | 0.548 | 0.634 | 0.652 | 0.633 | 4.761 | 4.701 | 4.793 | 4.769 | 4.395 | 4.81 | 4.739 | 4.575 | 4.693 | 4.515 | 4.597 | 4.558 |
| 0.668 | 0.78 | 0.674 | 0.75 | 0.548 | 0.695 | 0.648 | 0.764 | 0.603 | 0.763 | 0.584 | 0.94 | 4.366 | 4.618 | 4.609 | 4.694 | 4.703 | 4.637 | 4.887 | 4.709 | 4.81 | 4.378 | 4.468 | 4.36 |
| 0.569 | 0.647 | 0.723 | 0.574 | 0.684 | 0.6 | 0.651 | 0.601 | 0.744 | 0.654 | 0.646 | 0.701 | 4.709 | 4.771 | 4.758 | 4.551 | 4.772 | 4.835 | 4.637 | 4.756 | 4.781 | 4.717 | 4.679 | 4.797 |
| 0.529 | 0.673 | 0.562 | 0.568 | 0.499 | 0.629 | 0.633 | 0.665 | 0.57 | 0.663 | 0.595 | 0.669 | 5.063 | 4.606 | 5.092 | 4.594 | 4.72 | 4.42 | 4.776 | 4.593 | 4.584 | 4.496 | 4.647 | 4.676 |
| 0.6 | 0.57 | 0.59 | 0.604 | 0.592 | 0.638 | 0.621 | 0.651 | 0.594 | 0.641 | 0.562 | 0.564 | 4.689 | 4.805 | 5.041 | 4.591 | 4.895 | 4.527 | 4.71 | 4.734 | 4.653 | 4.469 | 4.563 | 4.697 |
| 0.595 | 0.778 | 0.584 | 0.707 | 0.554 | 0.763 | 0.61 | 0.87 | 0.6 | 0.651 | 0.594 | 0.728 | 4.939 | 4.726 | 4.762 | 4.73 | 4.83 | 4.644 | 4.783 | 4.826 | 4.579 | 4.489 | 4.577 | 4.597 |
| 0.63 | 0.553 | 0.561 | 0.594 | 0.587 | 0.613 | 0.603 | 0.621 | 0.579 | 0.564 | 0.593 | 0.618 | 4.812 | 5.158 | 4.654 | 4.772 | 4.726 | 4.88 | 4.598 | 4.712 | 4.497 | 4.613 | 4.684 | 4.569 |
| 0.504 | 0.586 | 0.594 | 0.584 | 0.562 | 0.591 | 0.613 | 0.584 | 0.551 | 0.628 | 0.664 | 0.625 | 4.916 | 4.895 | 4.949 | 4.993 | 4.845 | 4.811 | 4.709 | 4.81 | 4.845 | 4.468 | 4.575 | 4.626 |
| 0.565 | 0.775 | 0.627 | 0.598 | 0.603 | 0.954 | 0.622 | 0.64 | 0.626 | 0.65 | 0.612 | 0.591 | 4.823 | 4.548 | 4.998 | 5.047 | 4.609 | 4.972 | 4.638 | 4.657 | 4.556 | 4.615 | 4.519 | 4.65 |

A similar experiment was conducted on Hela cells, the Z’ value of which was determined to equal 0.694.

**Table. 1b.** The ratio of Green/Red signal of each well (HeLa cells) treated with 1XHBSS or 10 μM FCCP. Z’=0.694.

| 1X HBSS | | | | | | | | | | | | 10 μM FCCP | | | | | | | | | | | |
| --- | --- | --- | --- | --- | --- | --- | --- | --- | --- | --- | --- | --- | --- | --- | --- | --- | --- | --- | --- | --- | --- | --- | --- |
| 1.2 | 1.03 | 1.18 | 0.97 | 1.12 | 0.99 | 1.09 | 0.96 | 1.08 | 0.95 | 1.14 | 1.01 | 4.9 | 4.74 | 4.41 | 4.83 | 4.73 | 4.74 | 4.89 | 4.97 | 4.89 | 4.75 | 4.25 | 4.44 |
| 1.05 | 1.07 | 0.97 | 0.93 | 1.04 | 0.99 | 1.17 | 1.01 | 1.07 | 0.98 | 1.1 | 1.01 | 4.65 | 4.51 | 3.99 | 4.88 | 4.37 | 4.83 | 4.71 | 4.69 | 4.28 | 4.53 | 4.51 | 4.45 |
| 1.17 | 0.95 | 0.96 | 0.98 | 0.97 | 0.97 | 0.96 | 1 | 0.97 | 1 | 1 | 0.9 | 4.82 | 4.76 | 4.73 | 4.74 | 4.48 | 4.62 | 4.59 | 5.16 | 4.49 | 4.5 | 4.35 | 4.25 |
| 0.96 | 1.06 | 0.88 | 1.11 | 1.08 | 1.51 | 1.05 | 1.19 | 1.1 | 1.16 | 0.97 | 1.15 | 4.65 | 4.46 | 4.54 | 4.23 | 4.32 | 4.51 | 4.62 | 4.4 | 4.53 | 4.35 | 3.99 | 4.24 |
| 0.95 | 1.05 | 0.94 | 1.01 | 1.1 | 1.07 | 0.96 | 1.02 | 1.06 | 1.03 | 0.95 | 1.01 | 4.69 | 4.38 | 4.25 | 4.3 | 4.12 | 4.18 | 4.24 | 4.21 | 4.24 | 4.3 | 4.12 | 3.77 |
| 1.05 | 1 | 0.99 | 0.97 | 1 | 0.95 | 1.02 | 1.08 | 1.07 | 1.04 | 1.09 | 0.97 | 4.02 | 4.29 | 4.2 | 4.25 | 4.16 | 4.29 | 4.24 | 4.26 | 4.2 | 4.35 | 3.96 | 3.91 |
| 1 | 0.98 | 0.93 | 1.03 | 0.99 | 0.98 | 0.97 | 0.98 | 0.98 | 0.99 | 0.97 | 0.96 | 4.2 | 4.21 | 4.1 | 4.03 | 4.31 | 4.04 | 4.29 | 4.22 | 4.1 | 4.18 | 4.02 | 4.12 |
| 1.02 | 1 | 0.96 | 1.02 | 0.93 | 0.98 | 0.96 | 0.96 | 0.93 | 1.1 | 1.04 | 1.03 | 4.07 | 4.13 | 4.03 | 4.03 | 4.27 | 4.11 | 4.19 | 3.98 | 4.02 | 3.93 | 3.79 | 3.95 |
| 0.99 | 0.9 | 0.94 | 0.97 | 0.92 | 1.03 | 0.93 | 0.96 | 1.03 | 1.01 | 1.04 | 0.98 | 4.36 | 3.97 | 4.02 | 4.32 | 4.19 | 4.37 | 4.3 | 4.22 | 4.21 | 4.29 | 3.99 | 4.1 |
| 1.07 | 1.18 | 1.1 | 0.95 | 1.03 | 1.05 | 1.04 | 0.99 | 1.04 | 0.94 | 1.11 | 0.99 | 4.01 | 4 | 3.88 | 4.15 | 4.05 | 4.07 | 4.13 | 4.22 | 3.89 | 3.97 | 3.96 | 3.98 |
| 0.96 | 0.96 | 0.99 | 0.88 | 0.94 | 0.95 | 0.94 | 0.96 | 1.05 | 1.01 | 1.02 | 0.96 | 4.1 | 3.99 | 4.03 | 4.1 | 4.19 | 4.08 | 4.17 | 4.3 | 4.26 | 4.11 | 3.99 | 4.09 |
| 1.17 | 0.9 | 0.97 | 0.96 | 1.01 | 0.98 | 0.99 | 0.99 | 0.97 | 0.98 | 0.99 | 0.94 | 4.34 | 4.11 | 4 | 4.17 | 4.12 | 4 | 4.11 | 4.22 | 4.08 | 4.11 | 3.89 | 4.09 |
| 1.01 | 0.95 | 0.99 | 0.96 | 0.95 | 0.95 | 0.97 | 0.98 | 0.96 | 0.98 | 1.06 | 0.95 | 4.24 | 4.13 | 4.34 | 4.29 | 4.09 | 4.32 | 4.25 | 4.27 | 4.17 | 4.13 | 4.01 | 3.98 |
| 1.11 | 0.94 | 1.07 | 1.03 | 1.06 | 0.99 | 1.01 | 0.94 | 1.07 | 0.99 | 1.02 | 1.03 | 3.92 | 4.15 | 4.36 | 4.28 | 4.2 | 4.24 | 4.35 | 4.04 | 4.16 | 4.03 | 3.93 | 4.04 |
| 1.09 | 0.99 | 1.12 | 0.87 | 1.11 | 0.94 | 0.97 | 0.91 | 1.09 | 0.98 | 1 | 1.07 | 4.24 | 4.12 | 4.01 | 4.16 | 4.41 | 4.18 | 4.36 | 4.39 | 4.12 | 4.21 | 4.09 | 4.16 |
| 1.08 | 1.1 | 1.04 | 1 | 1.14 | 1.01 | 1.04 | 1.02 | 1.03 | 1.03 | 1.06 | 1.03 | 4.53 | 4.52 | 4.43 | 4.42 | 4.38 | 4.52 | 4.59 | 4.5 | 4.34 | 4.19 | 3.97 | 3.77 |

### 4. Toxic effects of m-MPI homogeneous assay kit on HepG2 and HeLa cells

To determine the potential toxic effects of m-MPI, cellular ATP levels in HeLa and HepG2 cells were measured at different time points after the cells were exposed to the dye using Dexorgen’s EnerCount ATP assay kit. The results indicate that the dye mixture does not affect the ATP levels in both HepG2 or HeLa cells (Fig. 4A and Fig. 4B).

**Fig. 4.**
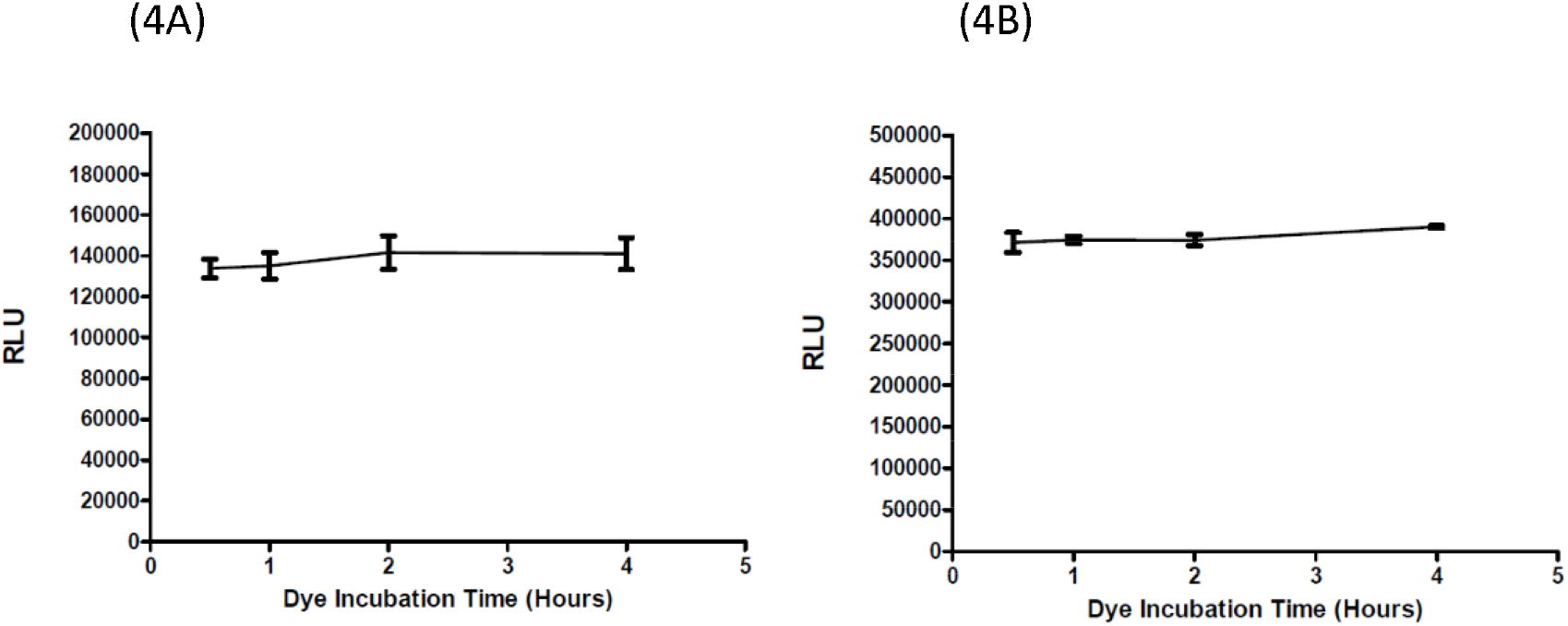
Relative ATP levels in HepG2 cells (4A) and in Hela cells (4B) after the cells were incubated with m-MPI over different amounts of time.

### 5. m-MPI assay on four different human primary cells

Four human primary cells—human hepatocytes, human adipocytes, human epidermal melanocytes, and human skeletal myoblasts—were cultured and plated on to 384-well black/clear plates as described in Materials and Methods. Four compounds: FCCP, Rotenone, Antimycin, and Valinomycin were tested as described in Materials and Methods. The m-MPI assay worked remarkably well on the different primary cells (Fig. 5).

**Fig. 5.**
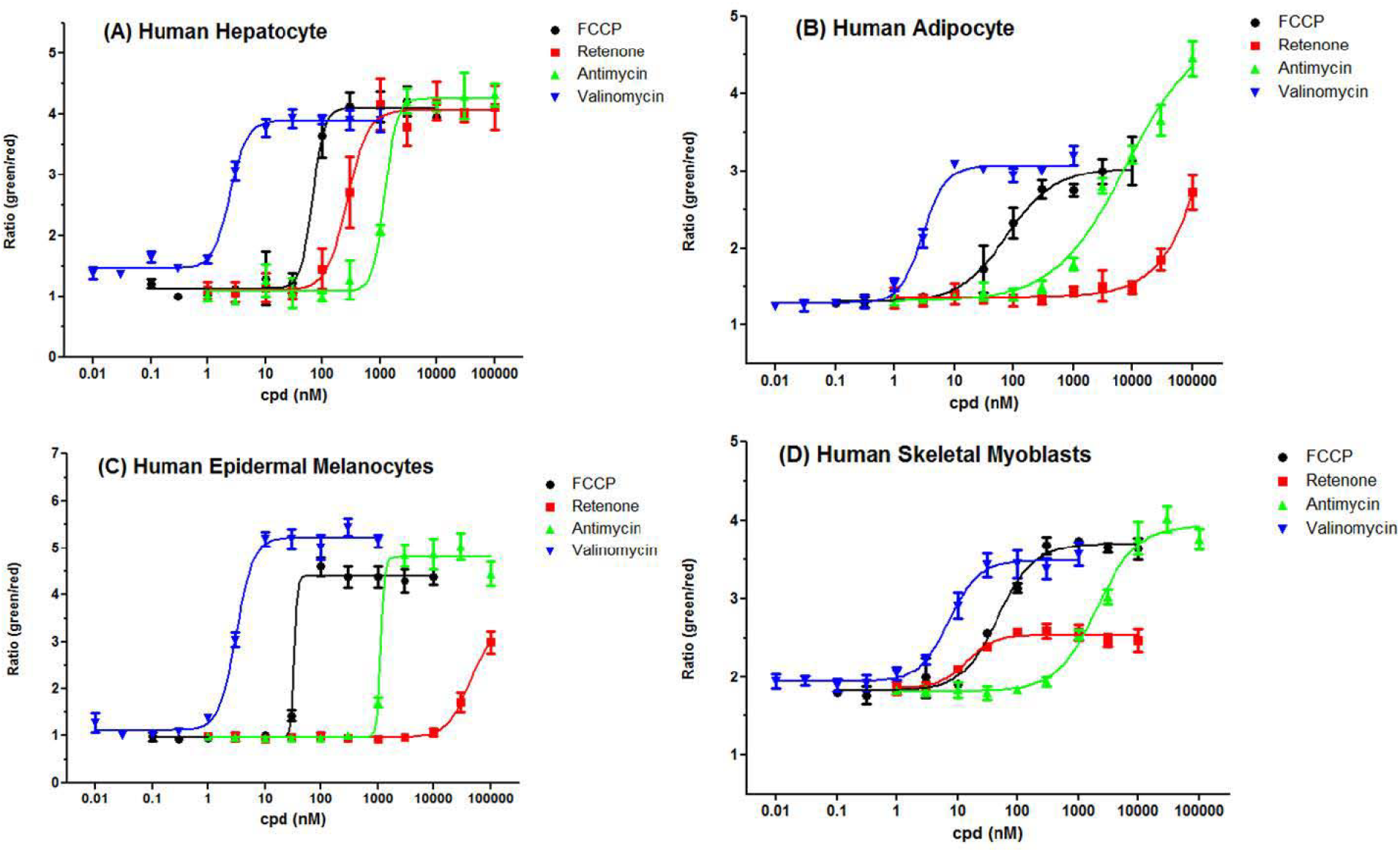
m-MPI Assay with different human primary cells. (A) Human hepatocytes, (B) Human adipocytes derived from the adult stem cells, (C) Human epidermal melanocytes cells, and (D) Human skeletal myoblasts cells

### 6. Test different compounds from LOPAC library on primary human hepatocytes

The LOPAC1280 library purchased from Sigma includes a biologically annotated collection of inhibitors, receptor ligands, pharma-developed tools, and approved drugs that impact most cellular signaling pathways and cover all major drug target classes.

The compounds from LOPAC 1280 library were picked and rearranged into a 96-well plate, as shown in the following table (Tab. 2a). Forty positive compounds and two negative compounds mentioned in the reference paper were chosen (13). Forty-seven samples not mentioned in the reference paper were randomly picked (supposed to be negative in HepG2 screening). FCCP was transferred into three wells as a positive control, and three wells of DMSO were used as negative control. Tyrphostin A9 (A9) was also used as a positive control. The assay was run as described in the Materials and Methods section with compound treatment for one or five hours. The final concentration of the testing compounds was 10 μM each.

Of the 41 positive compounds (other than FCCP), 28 compounds have more than 50% of FCCP activity with one hour of treatment. 24 compounds have more than 50% of FCCP activity with five hours of treatment. Three compounds have more than 50% of FCCP activity with five hours of treatment but less than 50% of FCCP activity with one hour of treatment. Therefore, a total of 30 compounds have more than 50% of FCCP activity in m-MPI assay. Among those 30 positive compounds, three compounds were negative with the HepG2 cell line but positive with primary hepatocytes, which are P7-A2, P8-F6, and P12-B11 in LOPAC library.

**Table. 2a:** Rearrangement of Compounds from LOPAC library to a 96-well compound-plate.

|  | 1 | 2 | 3 | 4 | 5 | 6 | 7 | 8 | 9 | 10 | 11 | 12 |
| --- | --- | --- | --- | --- | --- | --- | --- | --- | --- | --- | --- | --- |
| A | FCCP | P4-H08 | P7-H03 | P12-B09 | P14-G11 | P16-C02 | P2-G4 | P4-G3 | P7-A2 | P9-C4 | P12-A3 | P14-B10 |
| B | P1-G06 | P5-E08 | P8-G10 | P12-D06 | P15-A02 | P16-D02 | P2-E9 | P4-H11 | P7-F7 | P10-F12 | P12-G5 | P14-G4 |
| C | P2-F11 | P6-A02 | P9-E02 | P13-B10 | P15-B11 | P16-G02 | P2-B11 | P5-C5 | P7-C5 | P10-B2 | P12-B11 | P15-B7 |
| D | P2-G07 | P6-F04 | P9-H08 | FCCP | P15-C11 | P16-G03 | P3-C2 | P5-B11 | P8-D4 | DMSO | P13-E3 | P15-G2 |
| E | P3-H07 | P6-H04 | P10-F04 | P13-F08 | P15-D11 | A9 | P3-C8 | P5-F3 | P8-F6 | P10-G8 | P13-G10 | P15-B9 |
| F | P4-A02 | P7-C10 | P10-F05 | P14-B03 | P15-F06 | DMSO | P3-F9 | P6-C7 | P8-H4 | P11-G3 | P13-B6 | P16-A7 |
| G | P4-A03 | P7-C11 | P11-G07 | P14-C06 | P15-H08 | P1-G2 | P3-H2 | P6-F10 | P9-B9 | P11-B7 | FCCP | P16-G8 |
| H | P4-C10 | P7-D11 | P11-H08 | P14-G09 | P16-A10 | P1-C10 | P4-B7 | P6-H7 | P9-H2 | P11-H11 | P14-E3 | DMSO |
|  |  |  | Positive in HepG2 |  |  |  |  |  |  |  |  |  |
|  |  |  | Negative in HepG2 |  |  |  |  |  |  |  |  |  |
|  |  |  | Randomly Picked |  |  |  |  |  |  |  |  |  |

Individual compounds were then ordered from Sigma. The same batch of human hepatocytes from ZenBio was used for the subsequent assay. Eleven different concentrations of each compound were tested.

**Table. 2b:**
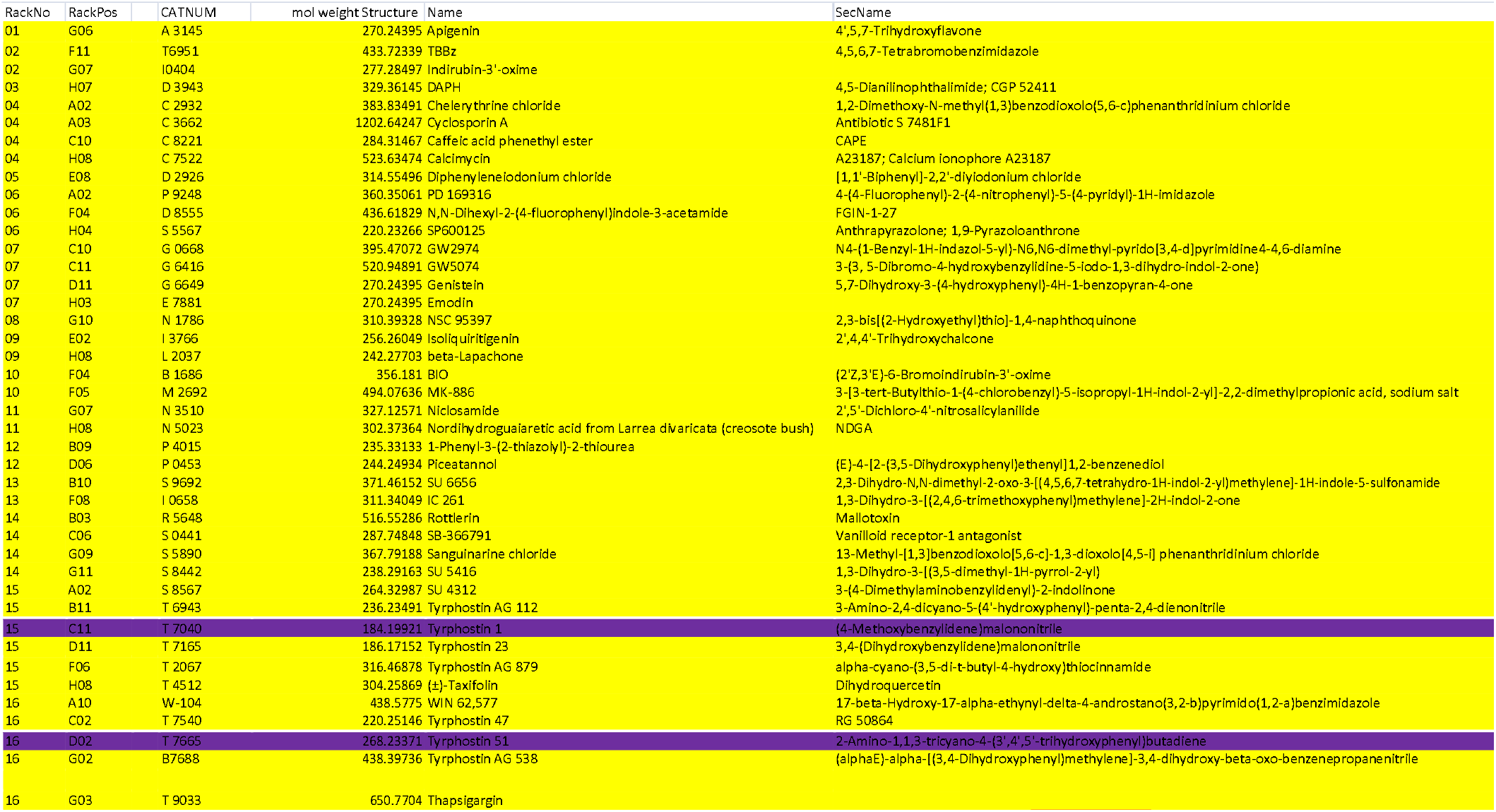
Positive and Negative Compounds from HepG2 Screening (Ref 12)

**Table. 2c:** Relative activity of the compounds compared with FCCP after one-hour treatment.

| Tab. 2c 1 h-Treatment |  |  |  |  |  |  |  |  |  |  |  |  |
| --- | --- | --- | --- | --- | --- | --- | --- | --- | --- | --- | --- | --- |
|  | 1 | 2 | 3 | 4 | 5 | 6 | 7 | 8 | 9 | 10 | 11 | 12 |
| A | 0.9828 | 0.9175 | 0.9765 | 0.975 | 0.4208 | 0.6292 | 0.1695 | 0.0139 | 0.5986 | 0.1747 | 0.095 | 0.1657 |
| B | 0.9174 | 0.8016 | 0.3959 | 0.8164 | 0.17 | 0.2342 | 0.2143 | 0.032 | -0.094 | 0.1183 | 0.0245 | 0.0716 |
| C | 0.9085 | 0.9976 | 0.977 | 0.2995 | 0.1252 | -0.099 | 0.0076 | -0.006 | 0.0703 | -0.05 | 0.7549 | -0.001 |
| D | 1.0025 | 0.122 | 0.1044 | 1.0043 | -0.126 | -0.096 | -0.037 | 0.1262 | 0.0789 | -0.18 | 0.2323 | -0.105 |
| E | 0.5264 | 0.7307 | 0.9842 | 0.136 | 0.4176 | 1.0017 | 0.1707 | 0.0585 | 0.7499 | 0.0411 | -0.114 | 0.0988 |
| F | 0.9978 | 0.5355 | 0.9896 | 1.0059 | 1.0205 | 0.0231 | 0.0307 | 0.037 | 0.081 | 0.006 | 0.1617 | 0.0129 |
| G | 0.0203 | 0.9804 | 1.0096 | 0.3151 | 0.1261 | -0.032 | 0.0882 | 0.1171 | -0.114 | 0.1693 | 1.0056 | 0.1094 |
| H | 0.551 | 0.897 | 0.6729 | 0.9838 | 0.0769 | 0.0757 | 0.1514 | -0.033 | 0.2381 | 0.0952 | 0.311 | 0.1606 |
|  |  | 0.5-0.85 |  |  |  | FCCP |  |  |  |  |  |  |
|  |  | >0.85 |  |  |  | <0.5 |  |  |  |  |  |  |

**Table. 2d:** Relative activity of the compounds compared with FCCP after five-hour treatment.

| Tab. 2d 5 h-Treatment |  |  |  |  |  |  |  |  |  |  |  |  |
| --- | --- | --- | --- | --- | --- | --- | --- | --- | --- | --- | --- | --- |
|  | 1 | 2 | 3 | 4 | 5 | 6 | 7 | 8 | 9 | 10 | 11 | 12 |
| A | 0.9828 | 0.8606 | 0.8744 | 0.9921 | 0.2209 | 0.3632 | 0.0891 | -0.073 | 0.6287 | 0.2079 | 0.0686 | 0.2911 |
| B | 0.0697 | 0.9597 | 0.6114 | 0.6996 | -0.163 | 0.4317 | -0.174 | -0.244 | 0.1085 | 0.1597 | 0.103 | -0.056 |
| C | 0.9182 | 0.8012 | 0.9332 | 0.1192 | -0.183 | -0.019 | -0.166 | 0.004 | 0.14 | 0.1485 | 0.7447 | 0.06 |
| D | 1.0134 | 0.8361 | 0.1884 | 1.0212 | -0.496 | 0.0728 | 0.0471 | -0.015 | 0.1359 | 0.1551 | 0.3918 | -0.215 |
| E | 0.3412 | 0.6505 | 1.0241 | 0.1967 | 0.3945 | 1.0174 | 0.1955 | 0.3523 | 0.5159 | 0.2344 | 0.0161 | 0.2049 |
| F | 0.9708 | 0.6826 | 0.9737 | 0.9707 | 1.0226 | -0.066 | 0.3974 | -0.073 | 0.2077 | 0.0007 | 0.1327 | -0.006 |
| G | 0.1373 | 0.9719 | 1.0122 | 0.3492 | 0.2045 | -0.008 | -0.097 | 0.166 | 0.1525 | 0.3192 | 0.9979 | 0.071 |
| H | 0.2797 | 0.4002 | 0.0929 | 0.9189 | 0.0555 | -0.105 | -0.035 | 0.1899 | 0.3141 | 0.1315 | 0.1413 | -0.09 |
|  |  | 0.5-0.85 |  |  |  | FCCP |  |  |  |  |  |  |
|  |  | >0.85 |  |  |  | <0.5 |  |  |  |  |  |  |

### 7. Confirmation of hits from LOPAC Library with human hepatocytes

The positive hits from LOPAC1280 Library were identified. Individual compounds were then purchased from Sigma. The same batch of human hepatocytes from ZenBio was used for the subsequent assay. The dose-response assays were performed (Fig. 6).

**Fig. 6.**
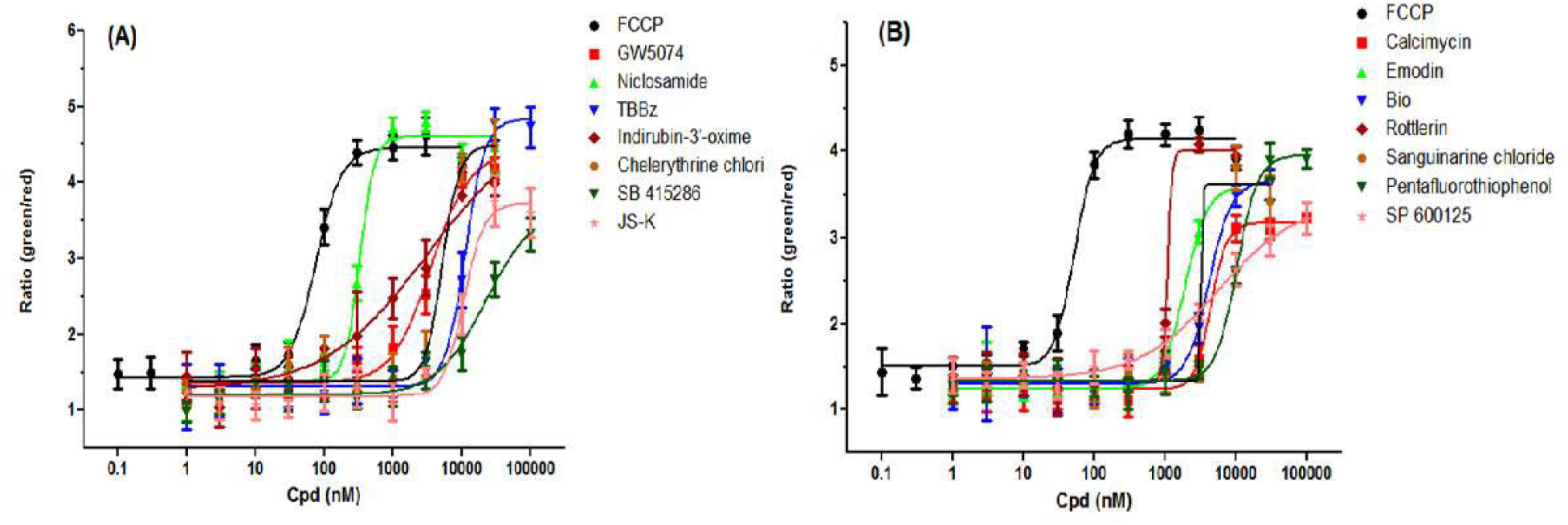
Confirmation of the hits from LOPAC Library with human hepatocytes.

## DISCUSSION

Newly developed or identified chemical compounds undergo toxicological analysis to determine if they are harmful to humans or other organisms. The degree of testing depends largely on the intended use of the compound. Drugs intended to treat human diseases undergo the most rigorous analysis through both in vitro and whole-animal testing. Whole-animal testing is the gold standard for these analyses because in vitro, it is impossible to replicate the complexity of drug interaction systems present in an animal. However, whole animal studies are extremely expensive, time-consuming, and often low throughput (14). Even extensive animal testing cannot provide a mechanistic understanding of toxicity (15). High costs preclude in vivo analysis of all newly synthesized chemicals in animals and limit the number of drug candidates that can be evaluated. Consequently, potential drug candidates usually go through a series of in vitro tests to screen out potentially toxic compounds before animal testing begins. However, growing ethical concerns about animal testing in combination with the relative cost-effectiveness of in vitro toxicity testing has intensified focus on in vitro testing.

Alternatives to whole-animal testing include assays on cell and tissue culture, use of tissue slices, toxicokinetic modeling and databases of known structure-toxicity relationships. These in vitro systems are ideally suited to investigate the molecular, cellular and physiological mechanisms of chemically induced toxicity that cannot readily be studied in vivo. Said systems are also useful when investigating known target organ and target species toxicity studies, as well as for answering specific questions about toxic effects. One justification for further developing in vitro toxicity tests is that they make toxicology a more scientifically based practice. In vitro testing allows considerably more control of experimental variables than whole-animal testing. Moreover, in vitro tests are less expensive and can determine toxicity more rapidly, especially with the progress of high-throughput (HTS) technologies.

Mitochondrial membrane potential (MMP) assays are a valuable tool used to evaluate drug toxicity in vitro. Such assays decrease the cost of toxicity screening and improve patient safety by screening out drugs with potential toxicity before they are tested on humans. Current assays are not ideal due to their usage of fluorescent cationic lipids not amenable to sensitive high throughput assays on all cell types, especially primary cells. As such, the newly developed fluorescent dye, m-MPI, will be used to develop a high throughput commercial assay that can be used with a multitude of primary cells and established cell lines.

The cell-based high-throughput assay described in this paper measures chemical toxicity using the novel dye, m-MPI, that was engineered to overcome many of the drawbacks of current dyes used to measure MMP. m-MPI rapidly partitions and moves to the cytoplasm or mitochondria; the resulting accumulation of dye in either location reflects the status of the MMP via fluorescence. JC-1, a parent dye of m-MPI that was previously used to measure MMP, functions by having its aggregates within the mitochondria fluoresce red and monomers in the cytoplasm fluoresce green (13). However, it is difficult to take advantage of these differential fluorescent properties due to the extremely low water solubility of JC-1. In comparison, the increased solubility properties of our dye, m-MPI, allows easy quantitative assessment of MMP status by simply calculating the ratio of green to red fluorescence. m-MPI can also be used with cells that do not work with JC-1 (such as primary cells). As such, we believe our m-MPI assay will more accurately assess hazards in a shorter time and at a lower cost.

We believe that the m-MPI assay can utilize human cells, including immortalized cells, primary cells, and stem cell-derived cells, and can function in a multi-well format with characteristics suitable for automated high-throughput screening.

Biologically, primary cells are a more accurate representation of mammalian cell biology than cultured cell lines. Thus, scientists are more likely to utilize primary cells rather than cultured cells in their studies. Primary cells are becoming increasingly prevalent in research studies, especially with new 3D cell culture technology which requires primary cells. We have tested the m-MPI assay kit on four human primary cells: human hepatocytes, human adipocytes derived from the adult stem cells, human epidermal melanocytes cells, and human skeletal myoblasts cells. The results indicate that the assay worked excellently on these primary cells.

iPSCs (induced pluripotent stem cells) are a type of pluripotent stem cell that can be generated directly from adult cells. They are a promising multipurpose research and clinical tool used to model diseases in the effort to develop and screen for drug candidates. Future studies could measure and compare cytotoxicity using the m-MPI assay kit on iPSC, iPSC-derived hepatocytes, iPSC-derived cardiomyocytes, and many more cell types.

In summary, we have developed a homogeneous assay based on m-MPI used in combination with a masking dye to measure the membrane potential change of mitochondria. This assay is nontoxic to the cells and has a Z’ value > 0.5 for HepG2 and HeLa cells. It is capable of measuring MMP change in human primary cells. From the data obtained, we believe that in the future, this m-MPI assay can be further miniaturized into a 1536-well format with human primary cells.

## ACKNOWLEDGMENTS

We thank Dr. Wei Zheng for critical reading of the manuscript. This research was supported by the National Institutes of Health. Contract number: HHSN271201300012C.

